# CRISPRpas: Programmable regulation of alternative polyadenylation by dCas9

**DOI:** 10.1101/2021.04.05.438492

**Authors:** Jihae Shin, Qingbao Ding, Erdene Baljinnyam, Ruijia Wang, Bin Tian

## Abstract

Well over half of human mRNA genes produce alternative polyadenylation (APA) isoforms that differ in mRNA metabolism due to 3’ UTR size changes or have variable coding potentials when coupled with alternative splicing. Aberrant APA is implicated in a growing number of human diseases. A programmable tool for APA regulation, hence, would be instrumental for understanding how APA events impact biological processes. Here, using a catalytically dead Cas9 (dCas9), we developed a method, named CRISPRpas, to alter cleavage and polyadenylation site (PAS) usage in 3’ UTRs or introns. We present key features that facilitate CRISPRpas, including targeting DNA strand, distance between PAS and targeting sequence, and strength of the PAS. For intronic PAS, we additionally analyze strengths of 5’ splice site and target location in intron. Our analyses implicate a dynamic competition between PAS usage and nascent RNA decay when RNA polymerase II elongation is blocked. We show modulation of APA of multiple endogenous genes including a gene that contains a single nucleotide polymorphism (SNP) that affects APA in the human population. CRISPRpas expands the CRISPR toolkit for perturbation of gene expression.

## INTRODUCTION

Cleavage and polyadenylation (CPA) is essential for 3’ end maturation of almost all eukaryotic mRNAs (1). CPA is carried out by the 3’ end processing machinery at the CPA site, commonly referred to as PAS. The strength or processing efficiency of a PAS is governed by its surrounding sequence motifs (2,3). While the upstream A[A/U]UAAA hexamer or other close variants are the most prominent motifs of PAS (4,5), other upstream and downstream motifs, such as UGUA, U-rich, and GU-rich motifs, additionally enhance PAS usage, often in a combinatorial manner (6–9). Mutations changing the PASs have been reported in many human diseases, such as thalassemia and systemic lupus erythematosus (10). Moreover, several recent studies have identified human single nucleotide polymorphisms (SNPs) near the PAS that can alter PAS usage (11,12).

Most mammalian genes have multiple PASs, resulting in alternative polyadenylation (APA) isoforms containing different coding sequences and/or 3’ untranslated regions (3’ UTRs) (13,14). APA in 3’ UTRs shortens or lengthens the 3’ UTR, thereby regulating aspects of mRNA metabolism, including stability, translation, and subcellular localization (15); APA sites in introns lead to transcripts encoding truncated proteins and can suppress gene expression (16,17). While most intronic polyadenylation (IPA) sites are generally considered “cryptic” and are repressed in normal conditions, emerging studies suggest that many developmental and pathological conditions lead to activation of IPA (16–20). While the biological importance of APA is increasingly appreciated, experimental strategies to modulate PAS usage are still limited.

The CRISPR/Cas9 system has emerged as a powerful tool for genome editing and gene regulation through transcriptional inhibition and/or activation (21–23). Cas9-mediated editing of PAS was employed in several recent studies to examine specific APA isoforms (18,24–26). However, genome editing permanently changes the DNA sequence and requires extensive manipulation of cells. A programmable APA at the RNA processing step is therefore desirable. Here we present a non-genomic editing method, named CRISPRpas, to alter APA. CRISPRpas delivers catalytically dead Cas9 (dCas9) (21,23) to the downstream region of a target PAS. Through blocking the progress of RNA polymerase II (Pol II), the dCas9 promotes usage of the PAS. We demonstrate effective APA isoform changes using reporter constructs and in multiple endogenous genes, including *ANKMY1*, whose APA is affected by an SNP in the human population. We elucidate several features that affect the efficacy of CRISPRpas, including targeting strand selection, distance from PAS to target locus, and PAS strength. When in the context of IPA, we further show the importance of splicing kinetics.

## MATERIAL AND METHODS

### Cell culture and transfection

Human HEK293T and HeLa Tet-On cells were cultured in high glucose DMEM with 10% FBS (Gibco) and 1% Penicillin/Streptomycin solution (Sigma). All cells were incubated at 37C with 5% CO2 and routinely checked by EVOS FL Auto Cell Imaging System (Thermo Fisher).

### Molecular cloning of plasmids

Information for plasmid construction is shown in the supplemental table S1.

### sgRNA design

sgRNA sequences were either designed using CRISPOR (27) which calculates gRNA specificity score as in a previous study (28) or were taken from previous publications. pGR9 plasmid was constructed to contain a Cas9 scaffold sequence. Oilgos were annealed and inserted into the pGR9 plasmid digested with BbsI. Oligonucleotides used for sgRNA cloning are listed in supplemental table S1. The 5’ and 3’ end 2’-O-Methyl and phosphorothioate modified sgRNAs were synthesized by Genscript (Piscataway, NJ).

### FACS analysis

48-72 hours after transfection with reporter plasmids, cells were collected by trypsinization. Green and red fluorescent signals were excited with 488nm and 561nm lasers using BD LSRFortessa X-20. Untransfected cells were used to determine background level. Signals were analyzed using BD digital software (DIVA) as well as home-made R codes. Briefly, cells were filtered by SSC and FSC. GFP-RFP double negative cells were also filtered. Log_2_ RFP and log_2_ RFP/GFP were calculated for each cell. Difference of mean log_2_ RFP/GFP was calculated by comparison between gene specific sgRNA and non-targeting control sgRNA. Standard of error of means (SEM) for Δ ratio was calculated by

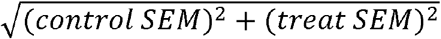

### PiggyBac stable cell line

To generate stable PiggyBac stable cell lines, we seeded cells in a twelve well plate. Next day they were transfected with HyPB7 and PiggyBac expression plasmids (System biosciences) using lipofectamine 3000 (Thermo Fisher) with 1ug total DNA. Cells were subsequently selected with 400ug/ml hygromycin for 6 days followed by monoclonal expansion. Successful genomic integration was confirmed by microscopy and western blot analysis.

### Transfection of sgRNAs for endogenous genes

HEK293T^dCas9^ cells were seeded in a twelve well plate one day before transfection. 1ug of pGR-sgRNA was transfected using Lipofectamine 3000 according to manufacturer’s protocol. Alternatively, 37.5pmole of synthetic sgRNA oligos were used. RNA samples were collected after 48 hours, and protein samples were collected after 72 hours.

### Transfection for reporter assay

200ng of reporter construct, 200ng of dCas9 encoding plasmid, 100ng of sgRNA encoding plasmid were mixed to transfect HeLa TetOn cells seeded in a 24 well plate using Lipofectamine 3000. Culture media was changed the next day, with 2ug/ml doxycycline. Induction of TRE promoter reporter constructs was carried out for 2 days. Alternatively, 200ng of reporter construct and 400ng of sgRNA-encoding plasmid were mixed to transfect HEK293T^dCas9^ cells.

### RT-qPCR

Total RNA was collected with TRIzol (Invitrogen). Residual genomic DNA was digested with TURBO DNase (Invitrogen) followed by inactivation of the enzyme. cDNA was synthesized from 2ug of total RNA using M-MLV reverse transcriptase (Promega) with an oligo(dT)18-25 primer. cDNA was mixed with gene-specific primers and subjected to RT-qPCR using Hot Start Taq-based Luna qPCR master mix (NEB). The reaction was run on a Bio-Rad CFX Real Time PCR system. Primers were designed to amplify specific APA isoforms, when needed. Primer sequences are listed in supplemental table S1. Two-tailed Student’s t-test was used to calculate significance of Δ Ct.

### Western blot

Protein concentration was determined using the DC Protein Assay (Bio-Rad). A total of 20ug of protein per sample was resolved using 4%-15% TGX stain-free gels (Bio-Rad), followed by immunoblotting using PCF11 (Proteintech, 23540-1) or GAPDH (CST, 5174) antibodies. Peroxidase donkey anti-rabbit IgG antibody was used as a secondary antibody (Jackson, 711-035-152). Signals on the blot was visualized by Clarity ECL reagent (Bio-Rad) with ChemiDoc Touch Imaging System. Signals were quantified using the ImageJ program and pre-stained molecular weight marker was used for protein size calculation.

### 3’READS+ library construction and sequencing

The 3’READS+ procedure was previously described (13,29). Briefly, 0.2-2ug of input RNA was captured using oligo(dT)25 magnetic beads and fragmented on beads with RNAse III. Partially digested poly(A)+ RNA fragments were ligated to 5’ adapter (5′-CCUUGGCACCCGAGAAUUCCANNNN) with T4 RNA ligase 1. The ligation products were incubated with biotin-T15-(T)5 LNA and digested by RNase H. Digested products were ligated to 3’ adapter with T4 RNA ligase 2. The final ligation products were reverse-transcribed and subjected for PCR amplification with index primers for multiplex sequencing. PCR products were size-selected with AMPure XP beads (Beckman) and subjected to quality control with ScreenTape (Agilent). Libraries were sequenced on llumina HiSeq (2 × 150 paired end reads). For 4sU labeling and fractionation, cells were cultures with 50uM of 4-thiouridine (Sigma) for 1 hours. 100ug of total RNA was biotinylated with biotin-HPDP, and subsequently captured by Streptavidin C1 Dynabeads (Thermo Fisher). The unbound flow-through (FT) fraction was also collected to represent pre-existing RNAs.

### RNA-Seq data analysis

GSE107648 sequencing data were re-analyzed to generate UCSC tracks for HEK293T cell expression of example genes. Raw reads from RNA-Seq were first mapped to the genome using STAR (v2.5.2) with default parameters (30) and converted to bigwig file format to create UCSC tracks.

### 3’READS analysis

3’READS data were processed and analyzed as previously described (31). Briefly, the sequence corresponding to 5’ adapter was first removed from raw 3’READS reads using Cutadapt (v1.18) (32). Reads with short inserts (<25 nt) were discarded. The retained reads were then mapped to the human genome (hg19) using bowtie2 (v2.2.9) (local mode) (33). The six random nucleotides at the 5’ end of reads 1 derived from the 3’ adapter were removed before mapping using the setting “-5 6” in bowtie2. Reads with a mapping quality score (MAPQ) <10 were discarded. Reads with ≥2 non-genomic 5’ Ts after alignment were called PASS reads. PASs within 24 nt from each other were clustered as previously described (13). The PAS reads mapped to genes were normalized by the median ratio method in DESeq (34).

### Stability score analysis

For all detected APA genes, the top two most expressed isoforms with PASs in the 3’ UTR were selected to calculate Stability Score of each isoform, which is log_2_ ratio of expression in the FT fraction over expression in the 4sU fraction. Δ Stability Scores dPAS versus pPAS for all APA genes detected in HEK293T cells are provided in supplemental table S2.

### Splice site strength

The strengths of 5’ and 3’ splice sites were calculated by the MaxEntScan program (http://hollywood.mit.edu/burgelab/maxent/Xmaxentscan_scoreseq.html) (35). Intron annotations were from RefSeq hg19 database.

### GTEx analysis

The raw RNA-seq data for GTEx data was downloaded from dbGaP (phs000424.v7) (36) including 5,032 RNA-seq samples. The SNP calling genotype data and phenotype data of GTEx were also downloaded from the same resource, only individuals covered by the RNA-seq were kept for the analysis. Gene expression and intronic APA analysis were done using Bioconductor package APAlyzer (v1.4.0) (37). Briefly, for gene expression analysis, reads mapped to CDS were used. For multi-exon genes, CDS region in the last exon was excluded. The read count was then normalized by CDS length and sequencing depth as transcript per kilobase million (TPM). For the IPA analysis, IPA sites with percent of samples with expression (PSE) > 5% were first extracted from PolyA_DB version 3.2 (38). The upstream and downstream regions of each IPA were then defined as the region between the site to its closest upstream 5’ or 3’ splicing sites (SS), and the region between the site to its closest downstream 3’SS. 5’SS and 3’SS information was obtained from RefSeq and Ensembl. The relative expression of IPA isoform (IPA-RE) = log_2_(a - b)/c, where a and b are read densities in IPA upstream and IPA downstream regions, respectively, and c is read density of the 3’ most exons. Only IPAs with at least five reads in each of the three regions were used for further analysis. For global APA analysis, IPA-RE of each gene were first standardized across samples, and then the median value of IPA-RE were used to represent the global IPA level for each tissue.

## RESULTS

### CRISPRpas alters PAS usage

CPA is known to be coupled with Pol II elongation (39). We thus hypothesized that a catalytically dead Cas9 (dCas9), which was previously shown to block Pol II elongation (23), might promote the usage of PAS. We named this approach CRISPRpas. We first tested CRISPRpas in the reporter system pRiG (40,41). A pRiG plasmid produces two APA isoforms due to the placement of two PASs in the vector (illustrated in Fig. 1A). Usage of its proximal PAS (pPAS) leads to expression of a short isoform (S) encoding RFP only, while usage of distal PAS (dPAS, from SV40 viral genome) leads to production of a long isoform (L) encoding both RFP and GFP. As such, fluorescence-activated cell sorting (FACS) analysis can be employed to interrogate the relative usage of the two PASs of the reporter vector.

**Figure 1.**
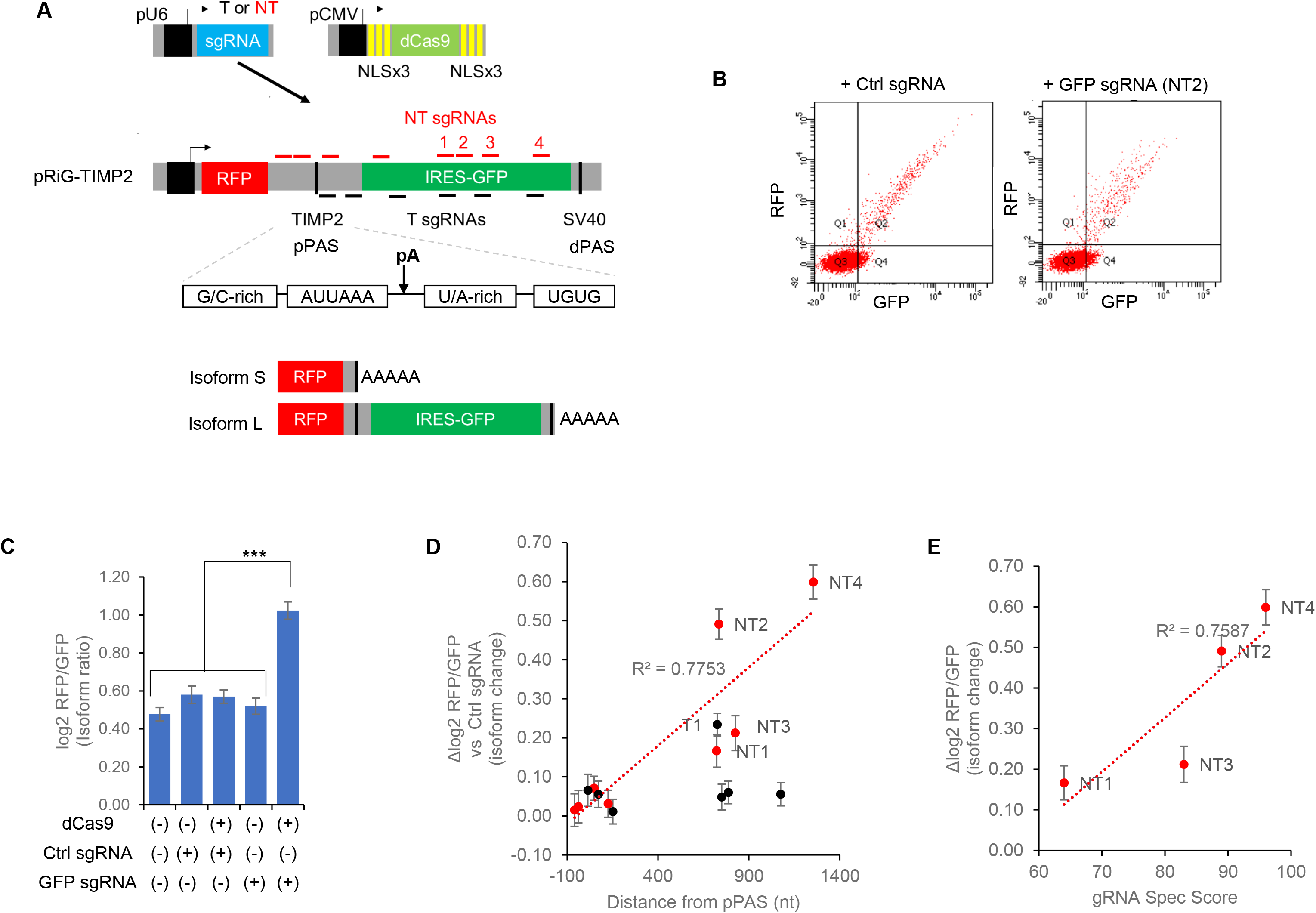
CRISPRpas alters 3’ UTR PAS usage in a reporter construct. A. Schematic of CRISPRpas using a reporter construct. U6 promoter drives expression of sgRNA targeting either the non-template (NT) or template (T) strand. sgRNA target locations are approximate. dCas9 was tagged with multiple copies of nuclear localization signal (NLS). Human *TIMP2* proximal PAS (pPAS) contains G/C rich, AUUAAA poly A signal, U/A rich and UGUG motifs. The pPAS and distal PAS (dPAS) are used by short isoform (S) or long isoform (L) respectively. B. Representative images of FACS data of pCMV-dCas9, pTRE-RiG-TIMP2 and sgRNA expressing vectors. Each dot indicates an individual cell, and X and Y axes indicate GFP and RFP intensity values, respectively. C. Comparison of mean log_2_ RFP/GFP of GFP sgRNA (NT2) with other samples. (+) and (−) denote presence and absence, respectively. For this and all other figures, error bar is SEM. * is p<0.05, ** p<0.01, and *** p<0.001 (Student’s t-test). D. Scatter plot for distance from pPAS versus Δ (compared to ctrl sgRNA) mean log_2_ RFP/GFP for eight NT and six T sgRNAs. Linear regression line is for NT sgRNAs. Four effective NT and one T sgRNAs are labeled on the plot. E. Scatter plot for gRNA specificity score and Δ log_2_ RFP/GFP for four NT sgRNAs.

We inserted a 180 nucleotide (nt) sequence spanning the pPAS of human *TIMP2* gene into the pRiG vector (Fig. 1A). The pRiG-TIMP2 plasmid was co-transfected into HeLa cells together with a plasmid encoding dCas9 with nuclear localization signals (NLSs) and a plasmid encoding sgRNA targeting to the GFP region that is located downstream of the pPAS. Compared to cells transfected with non-targeting control (Ctrl) sgRNA, those with GFP targeting sgRNA (NT2) showed decreased green fluorescence signal (Fig. 1B) indicating that the GFP-targeting sgRNA NT2 promoted the usage of pPAS. The observed effect of isoform change was not due to binding of sgRNA alone or non-specific activity of dCas9, because expression of neither dCas9 nor sgRNA alone had any effect (Fig. 1C).

We next tested several other sgRNAs targeting different loci on different strands (Fig. 1A). We found that none of the template strand (T) sgRNAs (black lines in Fig. 1A) had any effect on isoform changes, with the exception of T1, which showed a mild effect as measured by Δ log_2_ RFP/GFP (black dots, Fig. 1D). By contrast, four non-template (NT) sgRNAs (NT1 to 4) showed noticeable changes of APA (red dots, Fig. 1D). This DNA strand-specific regulation is in line with the fact that CRISPRpas works by blocking Pol II elongation (23). Interestingly, none of the sgRNAs whose target loci are close to pPAS (within 200 nt) elicited APA changes (Fig. 1D). The effectiveness of APA change and distance from the PAS were highly correlated for NT sgRNAs (r^2^=0.78, Fig. 1D) and the NT sgRNAs that are in the distance range from 770nt to 1301nt were effective (NT 1, 2, 3, and 4, Fig. 1D) for this reporter construct. For the four working NT sgRNAs, the effects on APA correlated well with their targeting efficiencies as predicted by sgRNA target specificity scores calculated by CRISPOR (27,28,42) (r^2^=0.76, Fig. 1E). Together, our results indicate that delivery of dCas9 and a sgRNA targeting non-template DNA can alter PAS usage and targeting region should not be close to the PAS.

Encouraged by our initial results, we next established a cell line HEK293T^dCas9^ with the dCas9 sequence inserted into the genome through the piggybac (PB) transposase system (Fig. 2A) for stable and robust expression of dCas9 in the cells. The dCas9 expression also drives BFP expression, enabling easy detection of dCas9 expression level (Fig. 2B). To investigate the effect of PAS sequence on CRISPRpas, we transfected HEK293T^dCas9^ with various pRiG plasmids containing different pPAS-flanking sequences (Fig. 2C). These PASs were derived from the human *CSTF3* gene (40,41) and had various strengths due to deletion mutations of surrounding sequences (Fig. 2C). The PAS strength is RiG-AE > RiG-AD > RiG-BD, and there was no PAS insertion in RiG (Fig. S2). Using the same sgRNA (NT2, Fig. 1A), we found that APA regulation by CRISPRpas (Δ log_2_ RFP/GFP, NT2 sgRNA vs ctrl sgRNA) was most effective in RiG-BD, then RiG-AD, and then RiG-AE, indicating that CRISPRpas works better when the PAS is weak (r^2^=0.88, Fig. 2D). Importantly, CRISPRpas did not function when there was no PAS (pRiG empty vector, Fig. 2C and 2D, orange dot). These results indicate that CRISPRpas specifically alters CPA when there is a PAS signal.

**Figure 2.**
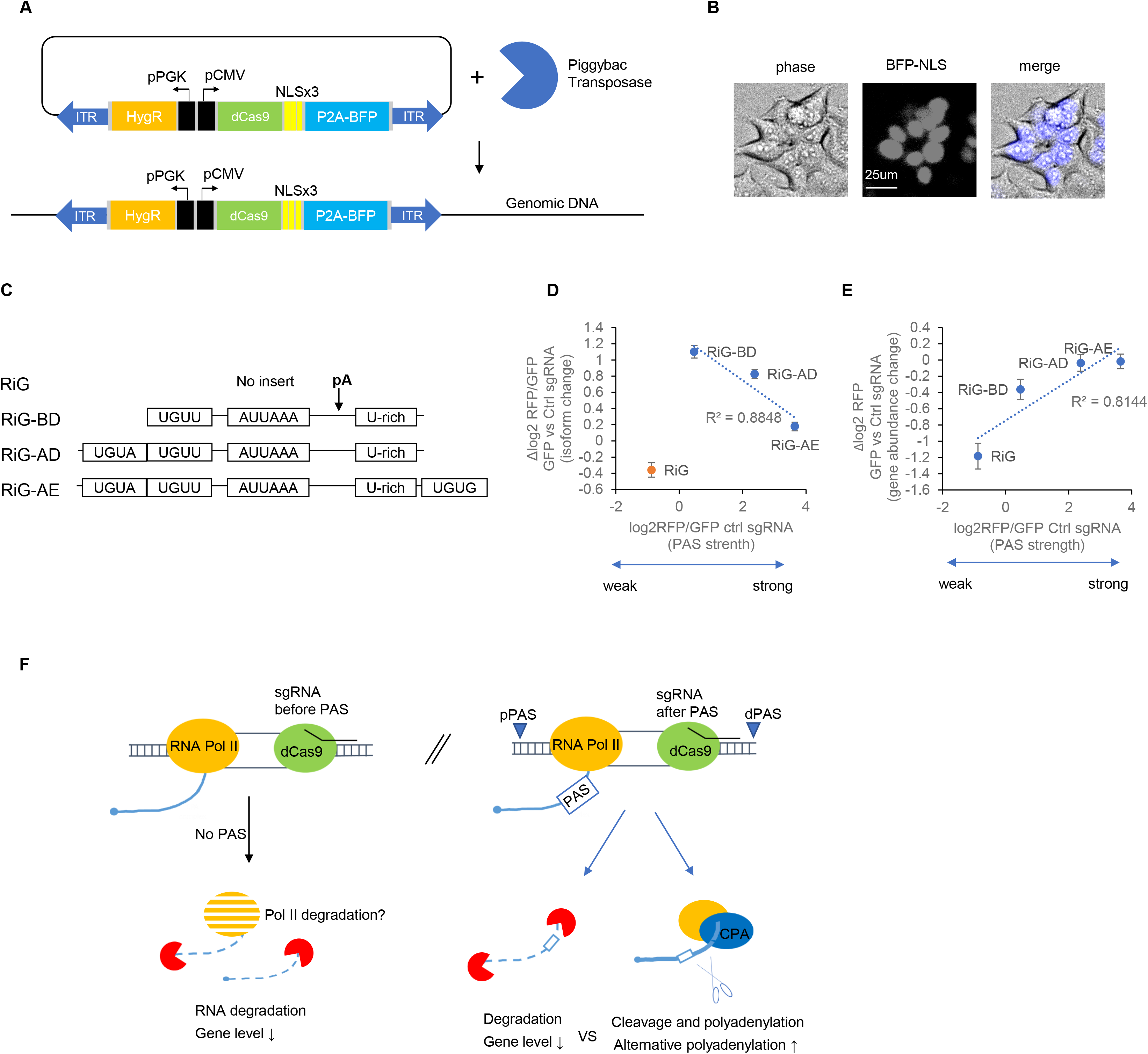
PAS strength affects CRISRPpas efficiency. A. Schematic for dCas9 expressing stable cell line created by the Piggybac transposase system. A bidirectional promoter system driving expression of dCas9-NLSx3-P2A (self-cleaving peptide)-BFP and Hygromycin resistant marker is flanked by transposon-specific inverted terminal repeats sequences for genomic integration. B. Nuclear BFP signals detected by fluorescence microscopy in HEK293T^dCas9^ cells. Phase, BFP and merged images are shown with a scale bar. C. Schematic diagram for pRiG constructs. White rectangles represent critical cis elements for PAS strength. pA stands for cleavage site. D. Scatter plot for PAS strength (basal log_2_ RFP/GFP) and isoform change which is Δ log_2_ RFP/GFP by GFP NT2 sgRNA compared to ctrl sgRNA in HEK293T^dCas9^ for pRiG vectors. Linear regression line is for three vectors with the pPAS (blue dots). pRiG has no PAS. E. Scatter plot for PAS strength (represented by log_2_ RFP/GFP) versus gene level abundance change (represented by Δ log_2_ RFP, test sgRNA vs ctrl sgRNA). Linear regression line is for all four vectors. F. A model for dCas9-mediated APA regulation. When the sgRNA target a region before the PAS, RNA Pol II is stalled without CPA, and the transcribing RNA gets degraded, presumably by the nuclear exosome. This leads to downregulation of gene expression. Pol II may also get degraded. Pre-mRNAs containing a weak PAS are more responsive to CRISPRpas, even though degradation is more apparent.

We next asked whether dCas9 would change the overall expression level of target gene. To this end, we measured the level of RFP signal as a proxy for gene expression. We found that RiG and RiG-BD decreased RFP expression in the presence of dCas9 (Wilcoxon test, p=3.8e-13 and 6.5e-4 respectively, Fig. 2E). There was no significant gene expression change for RiG-AD and RiG-AE (p>0.05, Fig. 2E). Interestingly, the level of decrease in RFP signal was inversely correlated with PAS strength: RFP signal of the pRiG was decreased the most, followed by RiG-BD, RiG-AD and RiG-AE (r^2^=0.81, Fig. 2E). A plausible explanation (depicted in Fig. 2F) is that while blocking Pol II elongation by dCas9 provides a time window for CPA, it also leads to degradation of nascent RNA, which is either associated with Pol II or has been evicted from Pol II. Transcripts with weak PAS are subjected to both degradation and CPA, whereas transcripts with strong PAS primarily to CPA.

### Regulation of endogenous 3’ UTR APA by CRISPRpas

The fact that the distance between PAS and targeting sequence is an important feature for CRISPRpas in our reporter system (Fig. 1D) indicates degradation of nascent transcripts can occur concurrently with CPA (Fig. 2F). To further explore this, we carried out CRISPRpas for an endogenous gene, *EIF1AD*. We designed three sgRNAs, targeting alternative (a) UTR region of *EIF1AD*. According to publicly available RNA-Seq data (43) and our own 3’ region extraction and deep sequencing (3’READS), a method specialized for interrogation of 3’ ends of polyadenylated RNAs (13), as well as our comprehensive PAS database (44), HEK293T cells express two major isoforms of this gene. We designed three sgRNAs targeting sequences 382 nt, 828 nt and 1045 nt away from the pPAS, aiming to test how distance would affect the efficacy of CRISPRpas (Fig. 1C). We detected isoforms by RT-qPCR using specific primers for common (c) UTR or aUTR regions. As we expected, we found that sgRNA distance from PAS was positively correlated with short isoform to long isoform ratio (the higher the ratio, the more 3’ UTR shortening, grey bars, Fig. 3B). Interestingly, suppression of gene expression (indicated by log_2_ fold change of cUTR region as compared to GAPDH mRNA) was observed when dCas9 was positioned close to the pPAS (sgRNA a, blue bar, Fig. 3B). These results indicate that distance between PAS and targeting sequence affect the outcome of CRISRPpas-regulated RNAs, again underlining the importance of distance between PAS and target site for efficient CRISRPpas.

**Figure 3.**
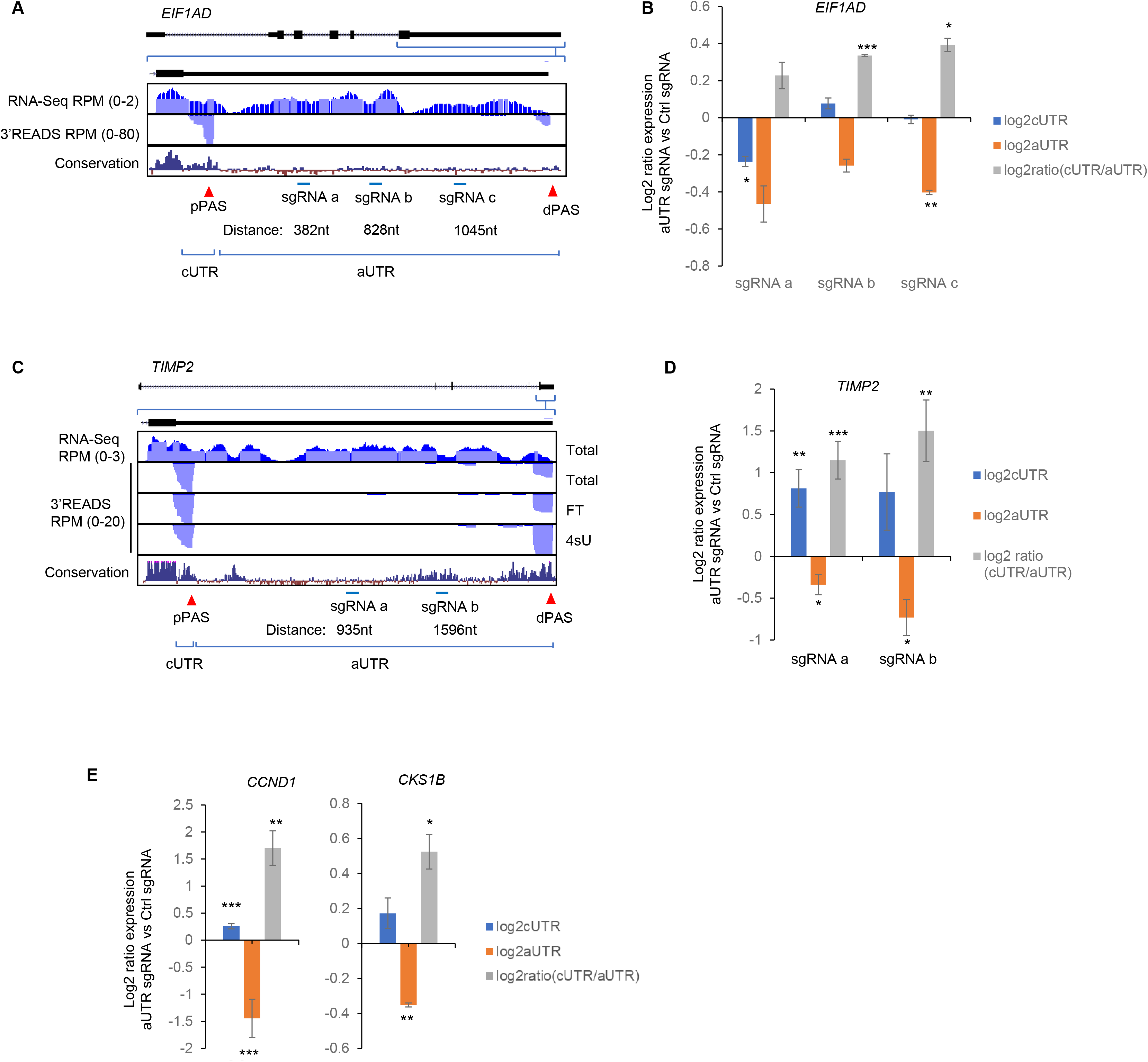
CRISPRpas modulates 3’ UTR APA of endogenous genes. A. Schematic of human *EIF1AD* gene structure. UCSC genome browser tracks for RNA-Seq and 3’READS are shown for the last exon of the gene. Sequence conservation based on 100 vertebrates is also shown. The two APA sites and three sgRNA targeting sites are also indicated, along with their distances from the pPAS. B. RT-qPCR analysis of relative amounts of short (blue) and long isoforms (orange) and ratio of short/long isoforms (grey). Plasmids encoding Ctrl or aUTR targeted sgRNAs were transfected in HEK293T^dCas9^ cells for 48 hours, and RNA was extracted for RT-qPCR. Log_2_ expression ratio was normalized to control. Error bars are standard error of mean of biological replicates (n=2). P value is calculated from Student’s t-test. (* is p<0.05, ** p<0.01, and *** p<0.001). C. Schematic of human *TIMP2* gene structure. UCSC genome browser tracks for RNA-Seq and 3’READS total, flow through (FT), and 4sU fractions are shown for the last exon of *TIMP2*. Sequence conservation based on 100 vertebrates is also shown. The two APA sites and common (c)UTR and alternative (a)UTR regions are indicated. The two sgRNA targeting sites are also indicated, along with their distances from the pPAS. D. Similar to Figure B for *TIMP2* gene (n=6, 4). E. Similar to Figure B, except that *CCND1* and *CKS1B* genes (n=3, 2) data are shown.

We also tested human *TIMP2* gene whose mouse homologue displayed dynamic APA regulation in differentiation (19). Its pPAS was also responsive to CRISPRpas in our reporter assay (Fig. 1). 3’READS data show that human cells express two major isoforms (Fig. 3C) as in mouse cells (29), and RNA-Seq data show robust expression of the aUTR region. We designed sgRNAs targeting aUTR region of *TIMP2* gene, 935 nt and 1596 nt away from pPAS. Using specific primers for cUTR or aUTR regions, we found that both sgRNAs (a and b) significantly decreased expression of aUTR region (orange bar) as well as increased short isoform to long isoform ratio (the higher the ratio, the more 3’ UTR shortening, grey bar, Fig. 3D). The sgRNAs also increased cUTR expression (~ 1.7 fold, Fig. 3D, blue bars), indicating a gene level upregulation. Because 3’ UTR-based mRNA stability has been reported in mouse cells (29,45), we hypothesized that *TIMP2* short isoform may be more stable than long isoform and such isoform-specific stability compensates the CRISPRpas-mediated gene downregulation. To investigate this possibility that more stable short isoform results in accumulated gene level upregulation, we metabolically labeled RNA with 4-thiouridine (4sU), and compared the 4sU-labeled, newly made transcripts to non-labeled, flow-through (FT) transcripts, which represent pre-exiting RNAs. In HEK293T cells, *TIMP2* long isoform is more enriched in 4sU fraction whereas short isoform using pPAS is enriched in FT fraction indicating higher stability of *TIMP2* short isoform (Fig. 3C, lower panels, Δ Stability Score dPAS vs pPAS = −0.24). Such increased mRNA stability of *TIMP2* suggests that CRISPRpas can be employed to manipulate gene expression. Similar results were obtained when we tested two additional endogenous genes, *CCND1* and *CKS1B*, (Fig. 3E).

sgRNAs can also be delivered to cells in the form of RNA. We next compared our initial plasmid-based sgRNA expression method to synthetic RNA oligo-based for CRISPRpas. For sgRNAs, both targeting sequence and Cas9 scaffold sequences were synthesized as RNA oligos with 5’ and 3’ end 2’-O-Methyl modifications (see Methods for detail). We found that synthetic sgRNAs worked better (by ~ 2 fold, Fig. S3) than plasmid-based, U6 promoter-driven sgRNA expression. Moreover, synthetic RNAs can regulate APA faster than plasmid-encoded sgRNAs within 24 hours post-transfection (Fig. S3). This result confirms our plasmid-based results, and additionally indicates the superiority of using synthetic sgRNAs in CRISPRpas.

### CRISPRpas regulates intronic polyadenylation

We next wanted to test CRISPRpas for IPA regulation, where CPA is coupled with alternative splicing (Fig. 4A). To this end, we first constructed a series of reporter plasmids based on the IPA site of *CSTF3* gene, a conserved site we previously found to be critical for *CSTF3* regulation (46). The construct series was named pRiniG (<u>R</u>FP <u>i</u>ntron <u>I</u>RES <u>G</u>FP) where the pPAS was flanked by 5’ and 3’ splice sites (SSs) of the intron 3 of human *CSTF3* gene. This reporter can measure efficiency of IPA because usage of pPAS leads to RFP expression whereas splicing and usage of dPAS leads to expression of both RFP and IRES-GFP (Fig. 4B).

**Figure 4.**
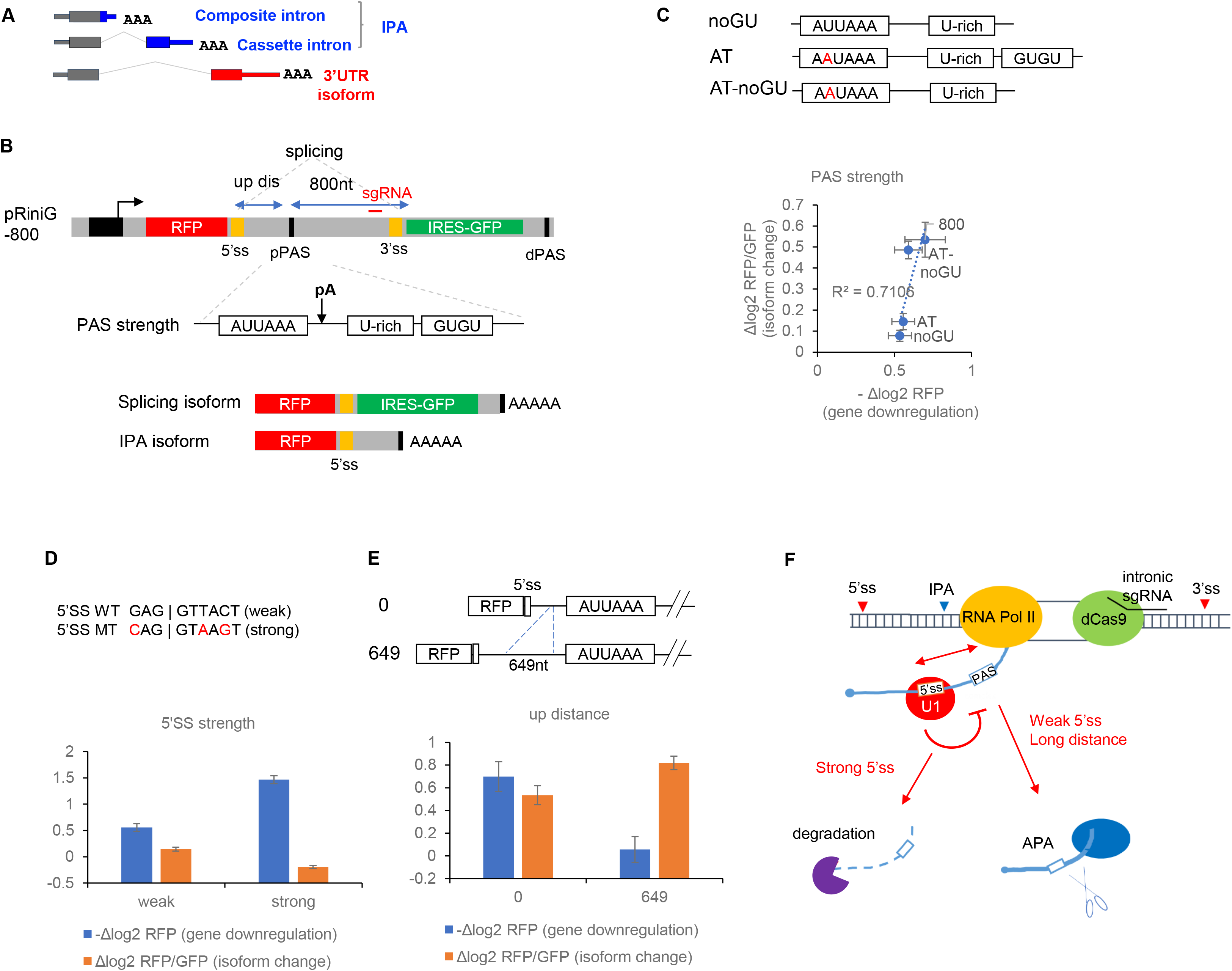
CRISPRpas regulates intronic polyadenylation in the reporter system. A. Schematic for different isoforms of transcripts including composite and cassette IPAs. B. Schematic of pRiniG-800 reporter construct to examine IPA regulation. Approximate location of the sgRNA is indicated. Regions containing 5’SS and 3’SS cloned from endogenous *CSTF3* gene are shown by yellow boxes. Splicing isoform is encoded by RFP-IRES-GFP via splicing. CPA cleaves at PAS and encodes IPA isoform. C. pRiniG-800 constructs were mutated to have various PAS strengths. In noGU construct, downstream GUGU sequence was deleted. In AT construct, weak wild type AUUAAA was mutated to strong AAUAAA sequence. AT-noGU have both mutations. Scatter plot for gene downregulation (−Δ log_2_ RFP) and isoform change measured by Δ log_2_ RFP/GFP (intronic sgRNA vs. ctrl sgRNA). Four reporters with different PAS strengths are shown. D. The wild type (WT) weak 5’SS sequence CAGGTTACT was mutated to a strong 5’SS CAGGTAAGT in pRiniG-800-AT. Bar graphs for gene downregulation (−Δ log_2_ RFP) and isoform change (Δ log_2_ RFP/GFP) using two constructs with different 5’SS strengths. Difference of the log_2_ ratio is comparison between intronic sgRNA and ctrl sgRNA. E. Distance between 5’SS to PAS was extended by inserting a 649 nt sequence. Bar graphs are the same as Figure D, except that data for different upstream distances are shown. F. Schematic model for CRISPRpas IPA regulation. A strong 5’SS inhibits CPA activity presumably by efficient U1 snRNP-mediated inhibition. This consequently subjects RNAs to degradation. When a weak PAS is effected on by CRISPRpas, upstream distance is a critical parameter of the CPA activity.

We first analyzed the effect of PAS strength (Fig. 4C). The parental pRiniG-800 was mutated to have various PAS strengths by mutating AUUAAA to stronger AAUAAA or deleting downstream GUGU sequences (Fig. 4C, S4A, and S4B). We noticed that medium strength PAS (800 and AT-noGU) are more sensitive to CRISPRpas than weak (noGU) or strong PAS (AT) (Fig. 4C), which is similar to our 3’ UTR data (Fig. 2D). This is because the strong PAS can trigger CPA regardless of dCas9, making it refractory to CRISPRpas. When the PAS is too weak, the RNA is not efficiently cleaved due to absence of a proper PAS signal. As shown in Fig. 4D, gene downregulation is correlated with isoform change, indicating dCas9 triggers CPA and degradation at the same time (r^2^=0.71).

We next tested the role of 5’SS strength by using the wild type (WT), a weak 5’SS (MaxEnt score at 1^st^-percentile) and a mutated, strong 5’SS (MaxEnt score at 96^th^-percentile) (46). As shown in Fig. 4D, strong 5’SS suppressed the IPA change (orange bar, Fig. 4D). This result is in agreement with the function of U1 snRNP in inhibiting CPA (47,48). Interestingly, the strong 5’SS also enhanced gene downregulation (blue bar, Fig. 4D), indicating that CPA suppression coupled with Pol II elongation blockade increases RNA degradation. Together, these results implicate a dynamic competition between PAS usage and nascent RNA decay when RNA polymerase II elongation is blocked.

We next hypothesized that perturbation of the distance between 5’SS and IPA site may have a similar effect to 5’SS strength change because extension of the distance promotes splicing and inhibits IPA (comparison between 800 and 649+800 in Fig. S4B). Intriguingly, increasing 5’SS-IPA distance enhanced IPA and mitigated degradation, similar to the effects of decreasing 5’SS strength (Fig. 4E). One possibility is that when splicing is inhibited due to RNA Pol II blockage before 3’SS, longer exposure of 5’SS may actually enhance CPA. This would be in line with a recent study showing the stimulatory model of U1-CPA complex (49), where U1 remodels and recruits CPA stimulating factors when U1:pre-mRNA base-paring is disrupted.

We also tested efficacy of CRISPRpas for reporters with long distance between PAS and 3’SS or weak 3’SS (Fig. S4C and Fig. S4D). In both cases, CRISPRpas was not found effective, due presumably to the high basal IPA activity of these constructs (Fig. S4B). Together, our data indicate that dCas9 can promote IPA when it targets a downstream region of IPA site, and highlight the importance of 5’SS in IPA regulation (Fig. 4F).

### Programmable APA by CRISPRpas changes gene expression

We and others recently identified a conserved IPA site in human and mouse *PCF11* genes (26,50). The IPA site is preceded by a weak 5’SS (MaxEnt score at 3^rd^-percentile in mouse and 5^th^-percentile in human). Our 3’READS data showed that *PCF11* had three prominent isoforms in HEK293T cells (Fig. 5A). We designed four synthetic sgRNAs targeting intron 1 of *PCF11* and transfected them into HEK293T^dCas9^ cells. Using primers designed to detect IPA isoforms or full-length (FL) isoforms, we found that three out of four sgRNAs significantly decreased full length (FL) expression and increased IPA/FL isoform ratio (Fig. 5B, except sgRNA-a). Interestingly, when the sgRNA target region was close to PAS (624 nt, sgPCF11-a), CRISPRpas was not efficient, despite high specificity scores for the sgRNAs. Indeed, a correlation could be discerned between distance from PAS and isoform change for *PCF11* gene (r^2^=0.51, Fig. 5C). These date are in agreement with the notion that the distance between PAS and targeting sequence is a critical factor for CRISPRpas. The decreased expression of the *PCF11* FL transcripts led to decreased protein production, which was detected by western blot analysis using anti-PCF11 antibody (~ 3.6 fold, Fig. 5D). These data highlight that we can effectively program endogenous IPA usage, which changes gene expression level.

**Figure 5.**
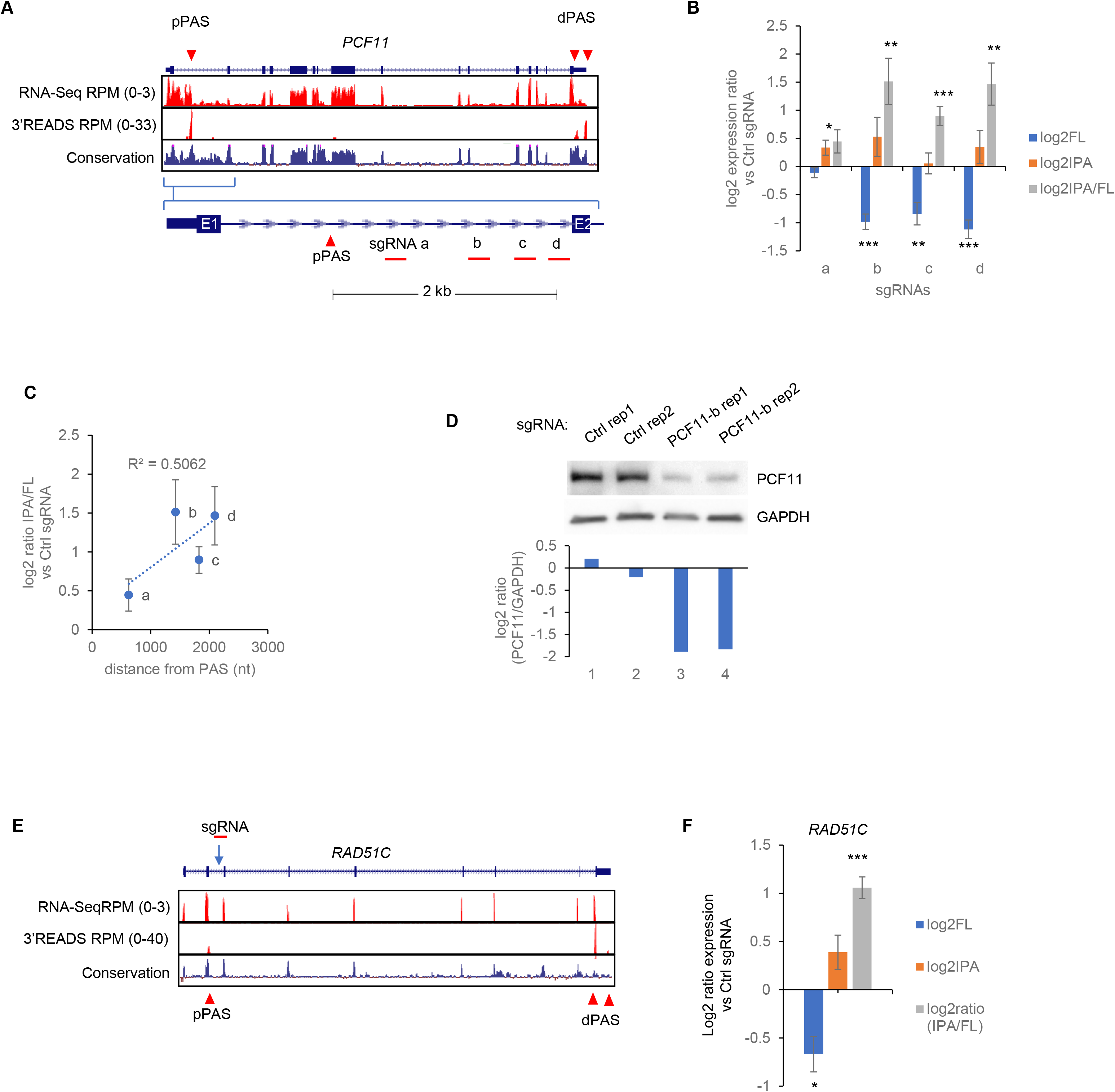
CRISPRpas alters intronic APA of endogenous genes. A. Schematic of human *PCF11* gene structure. Intronic and 3’ UTR PASs are indicated. Conservation track and expression data from RNA-Seq and 3’READS are also shown. Region between exon 1 and 2 is shown with approximate locations of four intronic sgRNAs. B. RT-qPCR analysis of relative amounts of FL (blue) and IPA isoforms (orange) and ratio of IPA/FL isoforms (grey). Ctrl and four *PCF11* synthetic sgRNAs were transfected in HEK293T^dCas9^ cells for 24 hours, and RNA was extracted for RT-qPCR. Log_2_ expression ratio was normalized to control. Error bars are from standard error of mean of biological replicates (n=3). C. Scatter plot for distance from PAS and log_2_ IPA/FL expression ratio indicating isoform change for four intronic sgRNAs with a linear regression line. D. Western blot analysis of Ctrl or PCF11-b sgRNA transfected HEK293T^dCas9^ cells. Protein lysates were collected 72 hours after transfection and band intensity was normalized to GAPDH using imageJ. E. Schematic of human *RAD51C* gene structure. Intronic and 3’ UTR PASs are indicated. Conservation track and expression data from RNA-Seq and 3’READS are also shown. Intronic sgRNA was designed in intron 2. F. RT-qPCR analysis of relative amounts of FL (blue) and IPA isoforms (orange) and ratio of IPA/FL isoforms (grey). Plasmids encoding ctrl and *RAD51C* sgRNAs were transfected in HEK293T^dCas9^ cells for 48 hours, and RNA was extracted for RT-qPCR. Log_2_ expression ratio was normalized to control. Error bars are from standard error of mean of biological replicates (n=3).

We additionally tested CRISPRpas on *RAD51C*, which is involved in DNA repair and mutations of which have been implicated in various cancers (51,52). Its IPA isoform, found to be regulated by termination factors (53), was clearly expressed in HEK293T cells (Fig. 5E). When we expressed a plasmid encoding sgRNA targeting intronic region downstream of IPA, we observed significant increase of the IPA/FL expression ratio, indicating efficient modulation of IPA by CRISPRpas for *RAD51C* gene (Fig. 5F).

### CRISPRpas can target a gene associated with APA-affecting SNPs

Recent studies have revealed many human SNPs associated with APA changes (11,12). Indeed, using our recently developed program APAlyzer program (54) and 4,126 Genotype-Tissue Expression (GTEx) RNA-seq datasets (55), corresponding to 27 tissues in 467 individuals, we identified 26,437 paQTLs associated with APA changes in human populations (see Methods for detail). We focused on the SNP rs13394744 (A>T), variants of which contained either the canonical PAS signal AATAAA or a weaker signal AATTAA. This SNP alters IPA in *ANKMY1*, as indicated by RNA-seq reads (Fig. 6A). Individuals with A/A homozygous alleles have higher expression of the IPA region than those with heterozygous A/T alleles or homozygous T/T alleles (Fig. 6B). Based on RNA-seq data of 17 human tissues, we detected an inverse correlation (*r*=-0.51, Pearson Correlation, Fig. 6C), indicating that increased IPA is associated with decreased gene expression. We designed an sgRNA targeting downstream region of *ANKMY1* IPA (Fig. 6A). Using HEK293T^dCas9^ cells, which had the A/A allele configuration, we were able to increase the IPA/FL isoform ratio and concomitantly decreased FL transcript expression (Fig. 6D). Taken together, our data on APA-affecting SNPs corroborate the notion that SNP-impacted APA can change gene expression and CRISPRpas can be used to alter naturally occurring APA differences.

**Figure 6.**
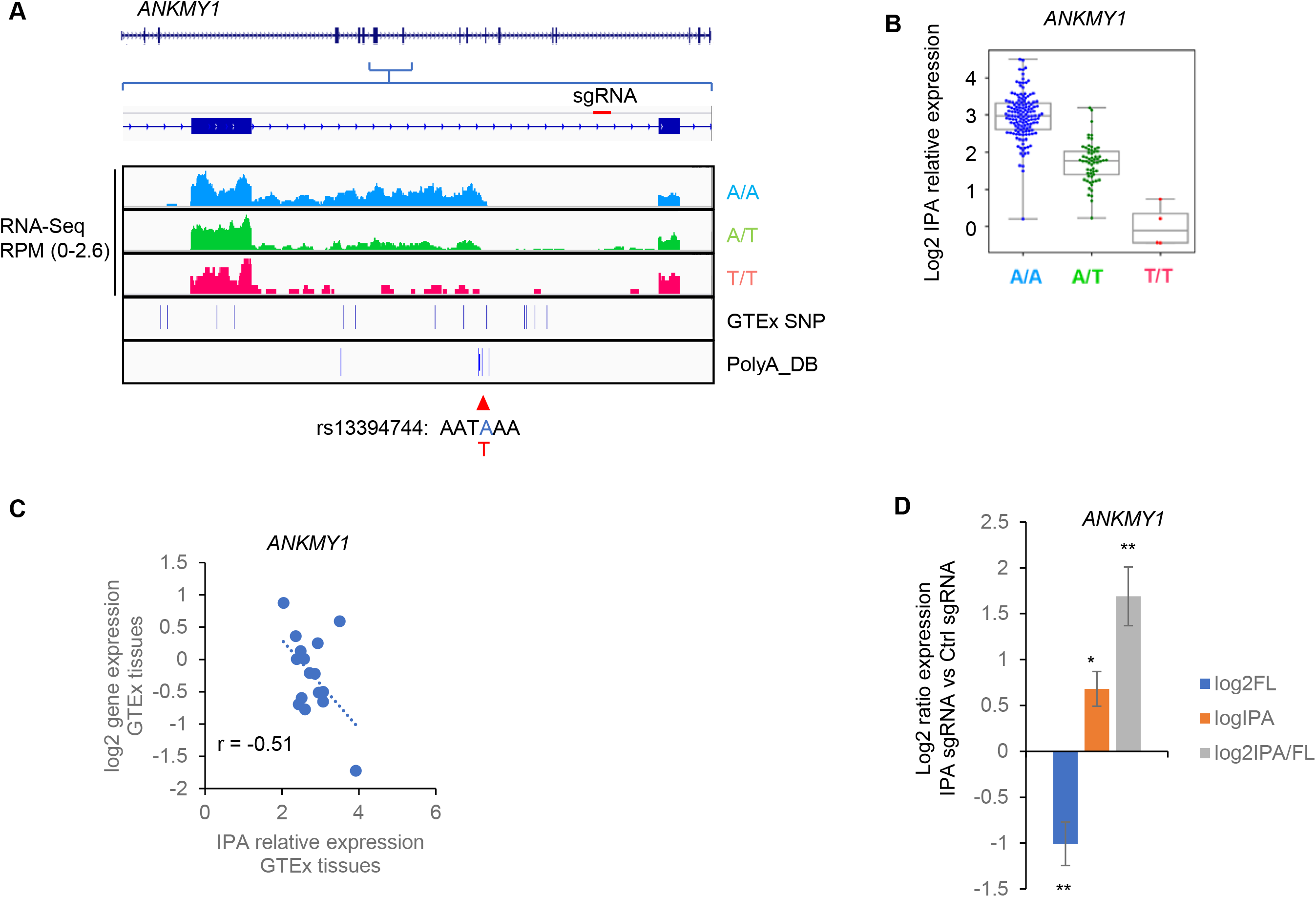
CRISPRpas alters intronic APA of *ANKMY1* gene. A. A SNP in human *ANKMY1* gene affects its APA. GTEx RNA-seq data samples are segregated by three allele types, as indicated. GTEx SNPs and PASs from polyA_DB3 are also shown. The SNP variants are AATAAA and AATTAA. Because AATAAA is stronger than AATTAA, intronic read coverage is lower in T/T samples than A/T or A/A samples. B. Relative expression (RE) of IPA of individuals with different alleles. C. Scatter plot for *ANKMY1* IPA RE vs normalized gene expression in 17 different tissues. D. RT-qPCR analysis of relative amounts of FL (blue) and IPA isoforms (orange) and ratio of IPA/FL isoforms (grey). Ctrl or sgRNAs targeting intron downstream of the pPAS of the *ANKMY1* gene were transfected in HEK293T^dCas9^ cells, and RNA was extracted for RT-qPCR. Log_2_ expression ratio was normalized to control. Error bars are from standard error of means of biological replicates (n=4). P value is calculated from Student’s t-test. (*, p<0.05; **, p<0.01; ***, p<0.001).

## DISCUSSION

In this study, we report a novel CRISPR/dCas9-mediated approach to regulate APA in human cells. We show that our CRISPRpas can induce change of mRNA isoforms with APA sites in 3’ UTRs as well as introns. Using reporter constructs, we found that PAS strength and the distance from PAS to targeting sequence are critical factors for the efficacy of CRISPRpas. Furthermore, we demonstrate the utility of CRISPRpas to modulate naturally occurring SNP-impacted APA differences between individuals.

CRISPRpas offers many advantages over other previously used methods for regulation of APA. Conventional Cas9-mediated gene editing of PAS has been used to manipulate APA of a gene, for example, addition of PAS to the end of coding region of *CCND1* gene (25) and deletion of *PCF11* intronic PAS (26,50). However, genome editing by Cas9 requires extensive manipulation of the cell and results in permanent changes to the genome. Several CRISPR methods have been developed to regulate RNA metabolism, such as RNA localization and alternative splicing (56–60), but not for APA manipulation. Our CRISPRpas method offers a programmable platform to regulate APA, complementary to currently available methods.

One potential shortcoming of CRISPRpas, however, is its effect on gene expression. This is because while Pol II stalling by physical collision with dCas9 increases the window of opportunity for CPA, it also leads to a greater chance of degradation of transcribing RNA. Hence, the distance between PAS and target site should be carefully examined when implementing CRISPRpas. Testing multiple sgRNAs would be necessary to identify the optimal one for both APA regulation and gene expression. In addition, consideration of isoform stability differences, as we did in this study, is also advisable. In this vein, we provide in supplemental table S2 isoform stability differences in HEK293T cells. Users of CRISPRpas can resort to this data for guidance, especially when HEK293T cells are to be used.

## DATA AVAILABILITY

Sequencing datasets generated in this study have been deposited into the GEO database under the accession number GSE161727. UCSC genome browser session can be accessed at https://genome.ucsc.edu/s/jihaesh/NAR_submission.

## ACKNOWLEDGMENTS

We thank members of BT lab for helpful discussions. We thank Dr. Renping Zhou (Rutgers school of pharmacy) for sharing reagents.

## Supplemental figure legend

**Figure S1.**
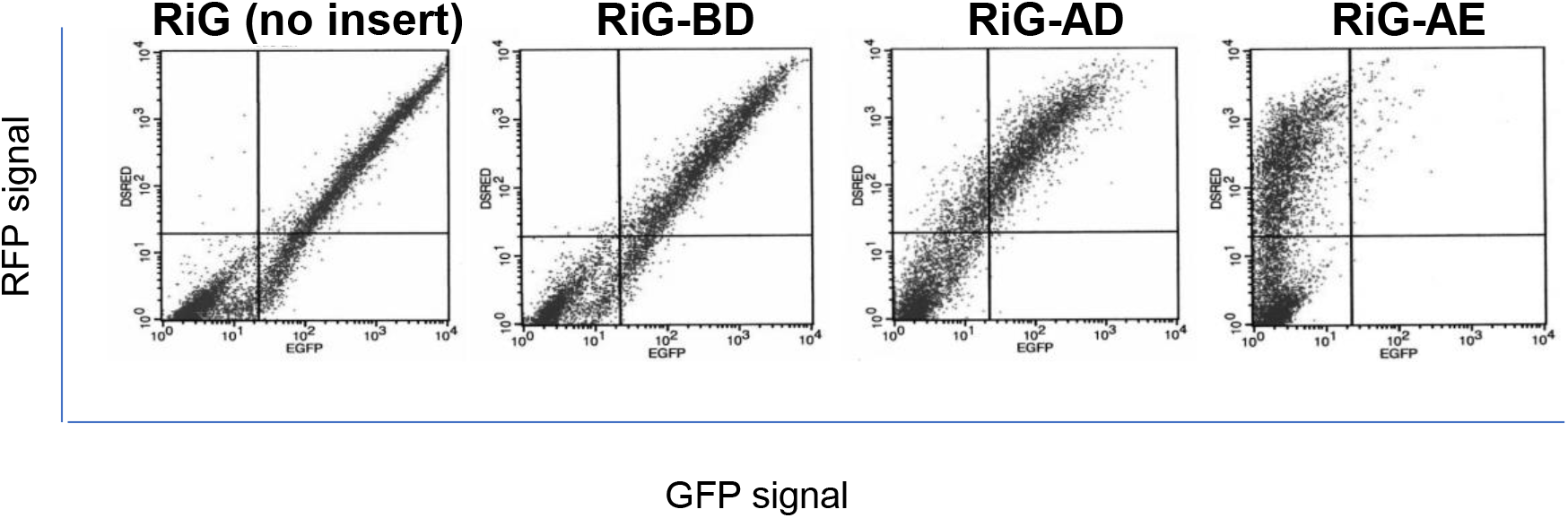
Representative images of FACS data of pCMV-RiG, pRiG-BD, pRiG-AD and pRiG-AE. Each dot indicates individual cell and X and Y axis indicate GFP and RFP intensity.

**Figure S2.**
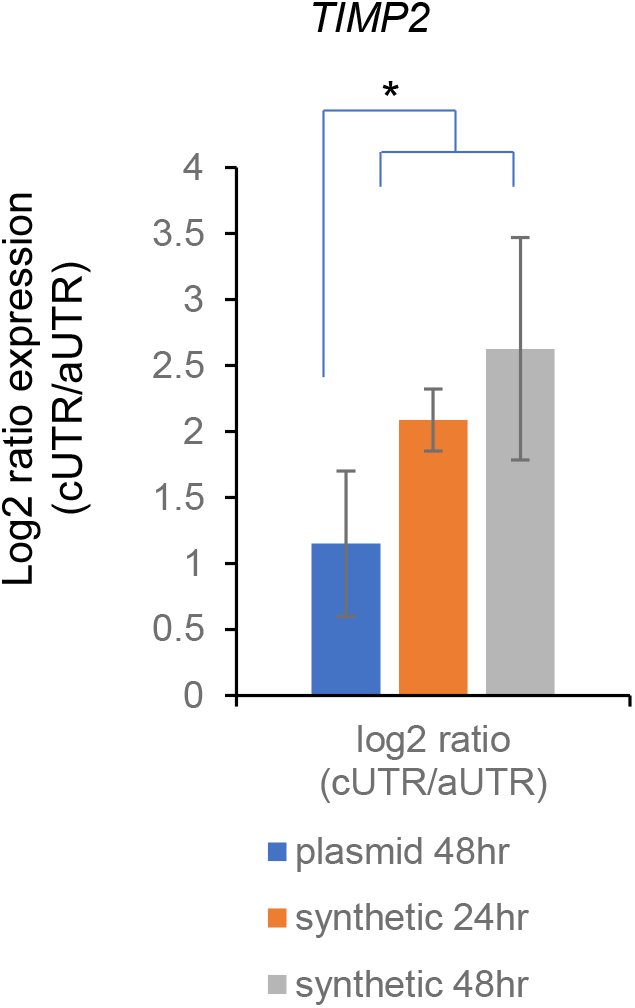
RT-qPCR analysis of ratio of TIMP2 short/long isoforms for plasmid based or synthesized sgRNAs. Plasmids encoding Ctrl or aUTR targeted sgRNAs were transfected in HEK293T^dCas9^ cells for 48 hours. Alternatively, chemically synthesized sgRNAs were transfected to cells for 24 or 48 hours. The RNA was extracted for RT-qPCR. Log2 expression ratio was normalized to control. Error bars are from standard error of mean of biological replicates (n=6,2,2). P value is calculated from Student’s t-test (*: p<0.05)

**Figure S3.**
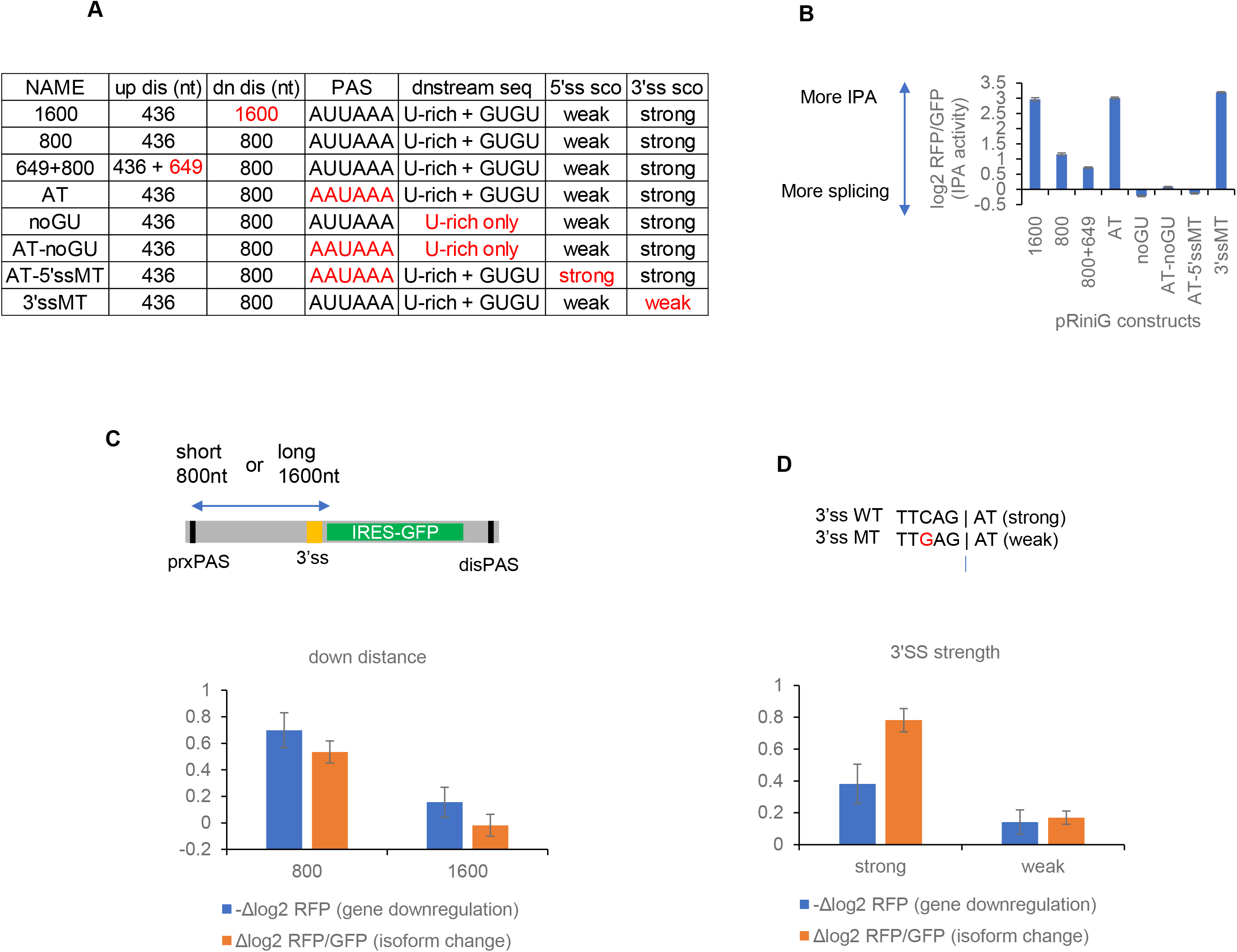
A. Table summarizing various pRiniG vectors used in this study. Difference from a parental vector pRiniG-800 was highlighted in red. B. Basal IPA activity measured by median log2 RFP/GFP ratio by FACS analysis for each construct is shown. Higher log2 RFP/GFP signal is associated with higher IPA activity and lower splicing activity. C. Distance between PAS and 3’SS was extended from 800 nt to 1600 nt. Bar graphs for gene downregulation (-delta log2 RFP) and isoform change (delta log2 RFP/GFP) using two constructs with different downstream distance. Difference of the log2 ratio is comparison between intronic sgRNA and ctrl sgRNA. D. Strong WT 3’SS TTCAGAT was mutated to weak 3’SS TTGAGAT. Bar graphs are same as Fig. S4C. In this case, difference of the log2 ratio is comparison between GFP sgRNA and ctrl sgRNA to target downstream of 3’SS.

**Supplemental table S1.**
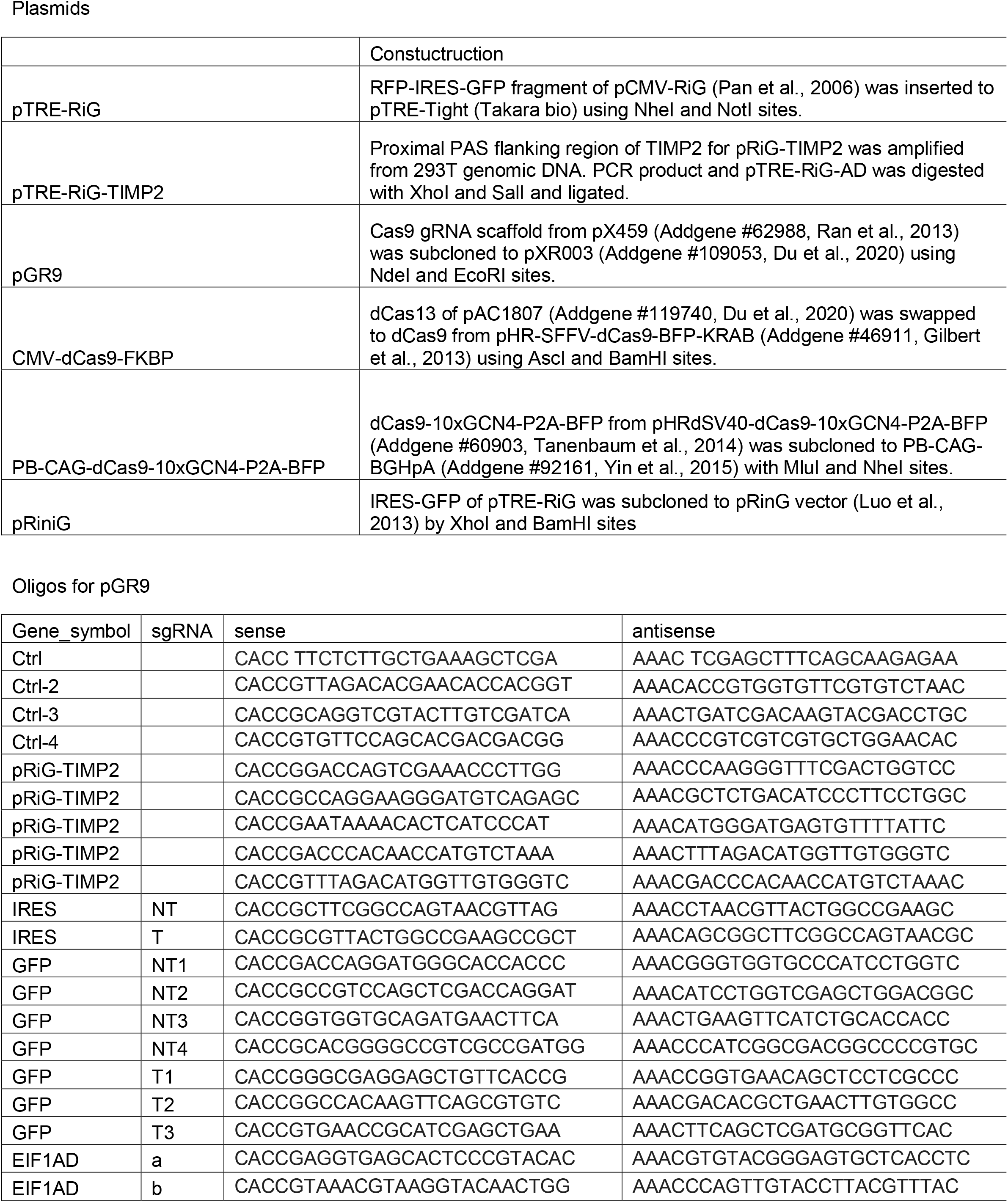
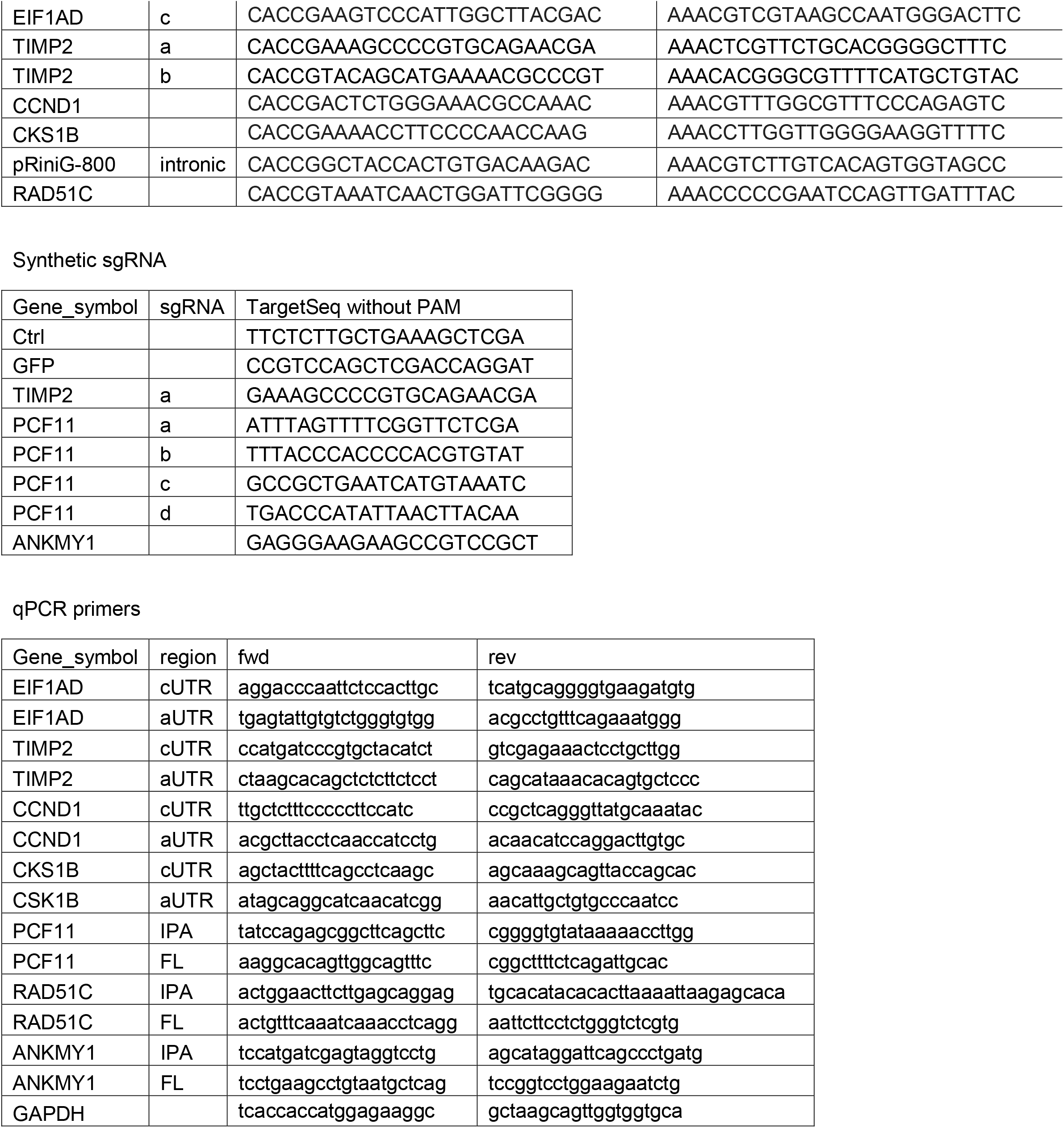

